# Mining Significant Features of Diabetes Mellitus Applying Decision Trees: A Case Study In Bangladesh

**DOI:** 10.1101/481994

**Authors:** Koushik Chandra Howlader, Md. Shahriare Satu, Avijit Barua, Mohammad Ali Moni

**Affiliations:** Dept of CSTE, Noakhali Science and Technology University, Bangladesh; Dept of CSE, Gono Bishwabidyalay; Faculty of Medicine, The University of Sydney, Australia

**Keywords:** Diabetes Mellitus, Feature Engineering, Data Mining, Decision Tree, Rule Extraction

## Abstract

Diabetes is a chronic condition which is associated with an abnormally high level of sugar in the blood. It is a lifelong disease that causes harmful effects in human life. The goal of this research is to predict the severity of diabetes and find out significant features of it. In this work, we gathered diabetes patients records from Noakhali Diabetes Association, Noakhali, Bangladesh. Thus, We preprocessed our raw dataset by replacing and removing missing and wrong records respectively. Thus, CDT, J48, NBTree and REPtree decision tree based classification techniques were used to analyze this dataset. After this analysis, we evaluated classification outcomes of these decision tree classifiers and found the best decision tree model from them. In this work, CDT unpruned tree shows highest accuracy, precision, recall, f-measure, second highest AUROC and lowest RMSE than other models. Then, we extracted possible rules and significant features from this model and plasma glucose, plasma glucose 2hr after glucose and HDL-cholesterol have been found as the most significant features to predict the severity of Diabetes Mellitus. We hope this work will be beneficial to build a predictive system and complementary tool for diabetes treatment in future.

## I. Introduction

Diabetes Mellitus (DM) is a driving cause of death and disability globally. It is a disorder of metabolism where the body uses digested food for energy [1]. It evolves when the body doesnt make sufficient insulin or is not capable to use insulin efficiently or both. There are existing three types of DM [2]. Type 1 DM results from the pancreas’s failure to produce enough insulin, Type 2 DM causes for insulin resistance, a state in which cells fail to respond insulin properly. Increased urine output, weight loss, excessive thirst, hunger, fatigue, yeast infections, skin problems, slow healing wounds and tingling or numbness in the feet are common symptoms of diabetes. Bangladesh has considered as highly diabetes population (adult) with 8.4% or 10 million according to research which published in WHO bulletin in 2013 [3]. Nearly half of the population with diabetes are not conscious about diabetes and don’t receive any treatment. A recent meta-analysis confirmed that prevalence of diabetes among adults had increased substantially from 4% in 1995 to 2000 and 5% from 2001 to 2005 to 9% from 2006 to 2010. According to the International Diabetes Federation (IDF), the prevalence will be 13% by 2030 [4]. The report also said that urban people were slightly more prone to diabetes than rural people. All these types of diabetes are dangerous and require treatment and if they are detected at the early stage, one can avoid different complications associated with them.

Data mining [5] has become popular area because it has powerful intensive and extensive applications. Analyzing complex, enormous biological data with various data mining techniques is an innovative and new field in biomedical sector. Matching and mapping strategies become so operative in diagnosis with data mining techniques [6]. Experts believe that data mining techniques in the healthcare industry will reduce the cost to 30% of overall healthcare spending. This goal of this study is to analyze diabetes dataset of patients and find out significant factors/conditions to happen diabetes. We collected N=220 patient’s data of diabetes from Noakhali Diabetes Association (NDA), Noakhali, Bangladesh which is affiliated with Bangladesh Diabetes Association (BDA) [3]. Then, our dataset was investigated with four decision tree (DT) based classifiers such as CDT, J48 [7], [8], NBTree and REPtree [10]. These classifiers were evaluated by different metrics such as Accuracy (Acc.), Kappa statistics (Ks.), Precision (Pr.), Recall (Rec.), F-measure (Fs.), Area Under Receiver Operating Characteristics (AUROC) and Root Mean Square Error (RMSE). Then, we identified the best classifier and extracted significant rules of diabetes from this model. We have also found several significant features that can help us to build decision support system (DSS) for physicians to predict the severity of diabetes.

This paper is divided into several parts. Several works which are related about diabetes research are discussed in section Section III describes step by step procedure how to we analyze diabetes data and mining significant features of it. Experimental outcomes are depicted and how to we extract significant features are described in section IV. Section V summarize our work with some limitations and denotes some future plan about diabetes research.

## II. Related Works

Different data mining techniques were developed various model for predicting diabetes. Sankaranarayanan et al. had used Frequent Pattern (FP) Growth and Apriori algorithm to generate association rules and found factors of diabetes [11]. When they required new rules of diabetes and these techniques could generate probable causes of diabetes in the form of association rule which could be used for fast and better clinical decision. In addition, Velu et al. represented Expectation Maximization (EM) Algorithm, H-Means Clustering (HMC), and Genetic Algorithm (GA) which were used to classify diabetes patient’s records [12]. These techniques were applied to generate individual clusters of similar symptoms. It was also claimed that HMC and double crossover genetic process based methods were shown better performance to compare other scales. Vijayan et al. showed that K-Nearest Neighbor (KNN), K-means, amalgam KNN and ANFIS were used to predict and diagnosis DM [13]. For maximizing expectation in a successive iteration cycle, they used EM algorithm for sampling diabetes data. They also used KNN algorithm for classifying objects and predicting them based on some closest training samples. Jianchao Han et al. preprocessed Prima Indian dataset by identifying and selecting attributes, removing outliers, normalizing data, visualizing data analysis, discovering hidden relationships and finally constructing a diabetes prediction model [14]. Bagdi et al. jointly used On Line Analytical Processing (OLAP) and Data Mining techniques to diagnosis diabetes by building a DSS that could response for complex cases [15]. They compared their evaluation metrics of this DT based algorithm with their proposed model. Kumari et al. proposed an intelligent and useful methodology to detect diabetes based on Artificial Neural Network (ANN) [16]. They also diagnosed their high dimensional medical data by support vector machine (SVM) and showed their result [17]. Aiswarya Iyer et al. classified diabetes data and extracted patterns employing naive bayes (NB) and DT [18]. Nahla et al. showed SVM as a promising tool to diagnosis DM [19].

## III. Dataset Description

We collected N=220 patients data with 13 attributes from Noakhali Diabetes Association (NDA), Maijdee, Noakhali, Bangladesh [3]. Within 13 attributes where 11 attributes are numeric and 2 attributes are nominal attributes. Different medical scales are inspected to identify this 13 attributes in this survey [20]–[27]. The details of diabetes data are represented in Table I. There were contained more missing values within 21 records in this dataset and removed them. Demographic characteristics of a person included age and gender are also considered the risk factors of diabetes.

**TABLE I.** Dataset Description

| S/N. | Attributes | Range | Relabeled values |
| --- | --- | --- | --- |
| 1 | Gender | Female, Male | gender |
| 2 | Age | 17-100 | age |
| 3 | Body Mass Index (kg/m<sup>2</sup>) | 17.31-38.19 | bmi |
| 4 | Sistolic Blood Pressure(mm Hg) | 60-180 | Systolic_press |
| 5 | Diastolic Blood Pressure (mm Hg) | 50-100 | diastolic_press |
| 6 | Plasma Glucose (mmol/l) | 3.43-23.68 | p_glucose |
| 7 | Plasma Glucose 2hr After Glucose(mmol/l) | 5.16-29.38 | glucose_two_hr |
| 8 | S Cholesterol (mg/dl) | 16-315 | s_cholesterol |
| 9 | HDL-Cholesterol (mg/dl) | 20-53 | hdl_cholesterol |
| 10 | LDL-Cholesterol (mg/dl) | 32-184 | ldl_cholesterol |
| 11 | S Triglyceride (mg/dl) | 66-589 | s_triglyceride |
| 12 | S Creatinine(mg/dl) | 0.6-3.34 | s_creatinine |
| 13 | Diagnosis | Healthy, Possible, Risky | class |

## IV. Methodology

In this study, there are considered some sequential steps to build a decision tree based classification model by analyzing diabetes data and predict the severity of diabetes. Those steps are described as follows 1:

- We have collected N=220 records of diabetes patient from NDA to predict the severity of this disease by analyzing these data. Then, several records and attributes were verified which have contained multiple unclear, duplicate and more missing values. All those missing values were filled manually with an approximation on individual class levels.
- There are considered 199 records of diabetes patient for further analysis. This data contained 52 healthy, 33 possible and 113 risky data of diabetes patient.
- There are considered four DT based classifiers such as CDT, J48, NBtree and REPTree to analyze diabetes patient data. We also analyze some DT based model which are applied with pruned and unpruned strategy. In this case, we compare their classification outcomes and find out the best classifier based on individual evaluation metrics. CDT(unpruned) shows the best performance than other algorithms.
- Then, we construct a DT of CDT(unpruned) and extract rules which is used to predict the severity of diabetes of individual people.

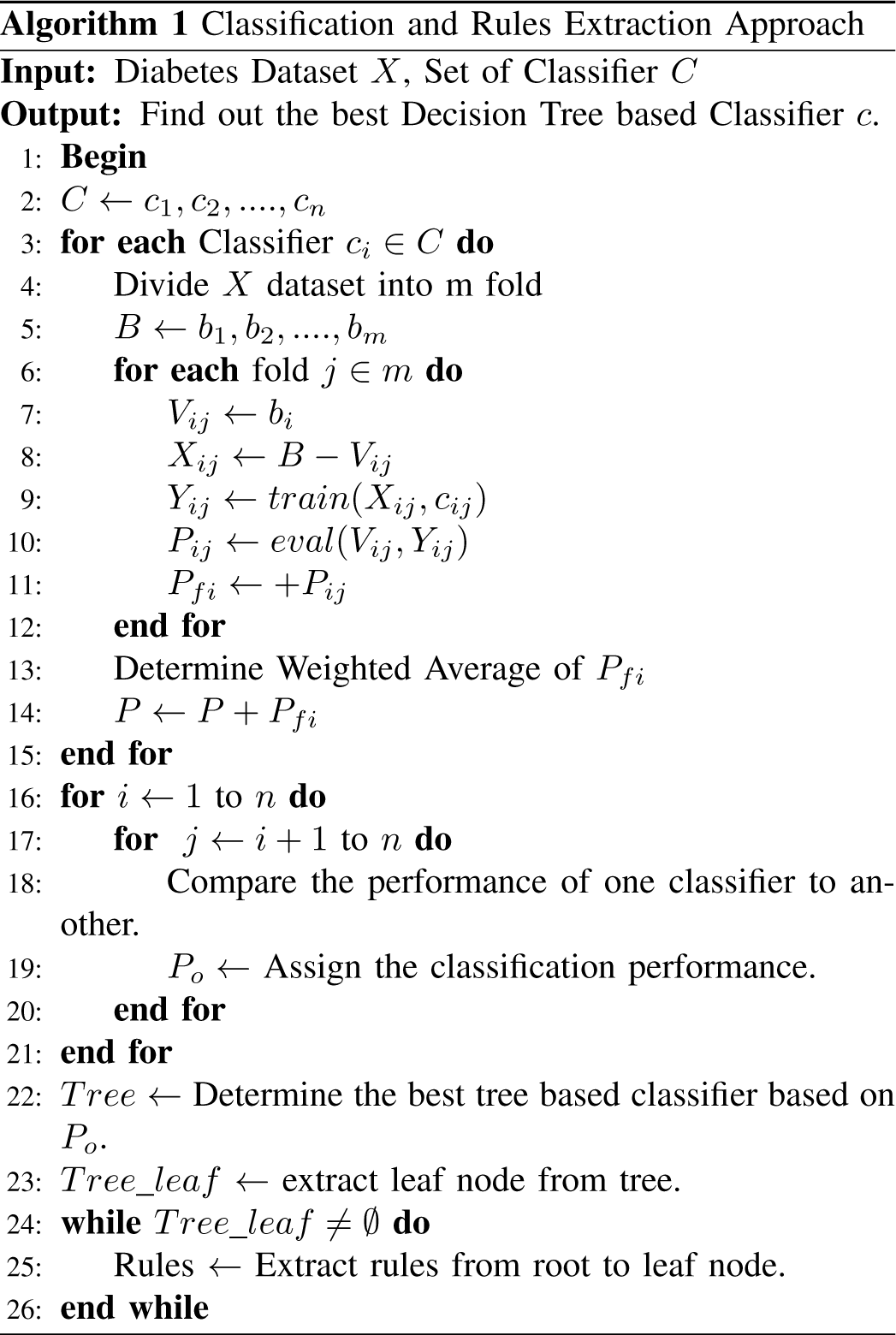

## V. Result & Discussion

In this experiment, we consider DT based classifiers to analyze this diabetes’s data in Weka [28]. We use 199 preprocessed instances of diabetes patients. There are used CDT (pruned), CDT (unpruned), J48 (pruned), J48 (unpruned), NBTree and REPTree to construct decision tree and find out significant rules of diabetes. Accuracy (Acc.), Kappa statistics (Ks.), and weighed average of Precision (Pr.), Recall (Rec.), F-measure (Fs.), AUROC and RMSE are used to find out the best DT based model. Table II shows classification outcomes of this classifiers.

**TABLE II.** Experimental Outcomes

| Classifier | Acc. | Ks. | Pr. | Rec. | Fs. | AUROC |
| --- | --- | --- | --- | --- | --- | --- |
| CDT(Pruned) | 96.48% | 0.939 | 0.965 | 0.965 | 0.964 | 0.971 |
| CDT(Unpruned) | 96.98% | 0.948 | 0.970 | 0.970 | 0.970 | 0.981 |
| J48(Pruned) | 96.48% | 0.939 | 0.965 | 0.965 | 0.965 | 0.978 |
| J48(Unpruned) | 96.48% | 0.939 | 0.965 | 0.965 | 0.965 | 0.978 |
| NBTree | 96.48% | 0.939 | 0.965 | 0.965 | 0.965 | 0.995 |
| REPTree | 96.48% | 0.939 | 0.965 | 0.965 | 0.964 | 0.971 |

In table II, CDT (unpurned) shows 96.78% accuracy where CDT(pruned), J48 (pruned), J48 (unpruned), NBTree and REPTree shows 96.48% accuracy. The value of kappa for CDT(unpurned) is 0.948 where CDT(pruned), J48 (pruned), J48 (unpruned), NBTree and REPTree shows 0.939. Besides, precision, recall and f-measure are found for CDT(unpruned) are 0.970, 0.970 and 0.970 respectively. On the other hand, CDT(pruned), J48 (pruned), J48 (unpruned), NBTree and REPTree show 0.965 for precision and recall respectively. J48 (pruned), J48 (unpruned) and NBTree show 0.965 and CDT(pruned) and REPTree show 0.964 for f-measure respectively.

Accuracy, Kappa statistics, precision, recall and f-measure shows biased outcomes in this experiment. So, we also consider another two metrics which is AUROC and RMSE. NBTree shows the highest AUROC (0.995) but it shows worse outcome than other classifiers. CDT(unpruned) shows second highest AUROC than others. Fig 1 shows AUROC of different DT based models. When we consider RMSE, CDT(pruned) and REPTree show 0.1507, J48(pruned) and J48(unpruned) show 0.1552, NBTree shows 0.1475 residuals. But, CDT(unpruned) shows lowest RMSE than other classifiers. So, CDT(unpruned) is considered as the best DT based model for extracting significant rules from it. CDT(unpruned), J48(pruned) and J48(unpruned) are build almost same DT. Fig 2, shows the DT which is given as follow:

**Fig. 1.**
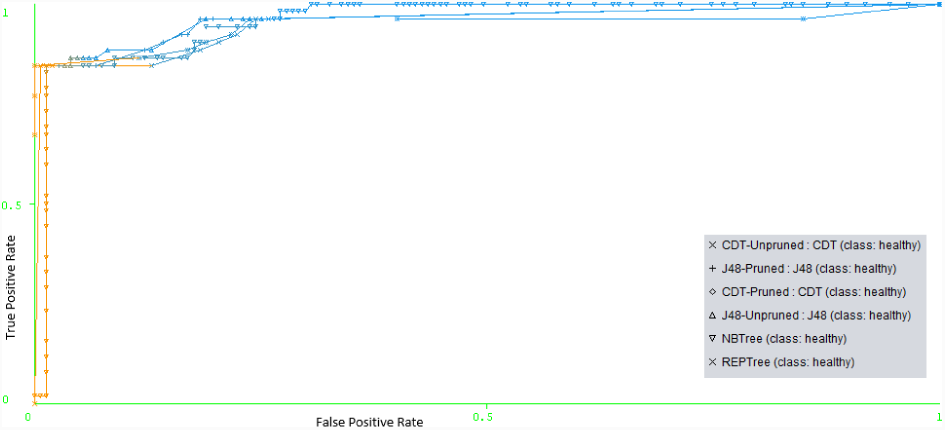
ROC Curve of different Tree Based Classifiers.

**Fig. 2.**
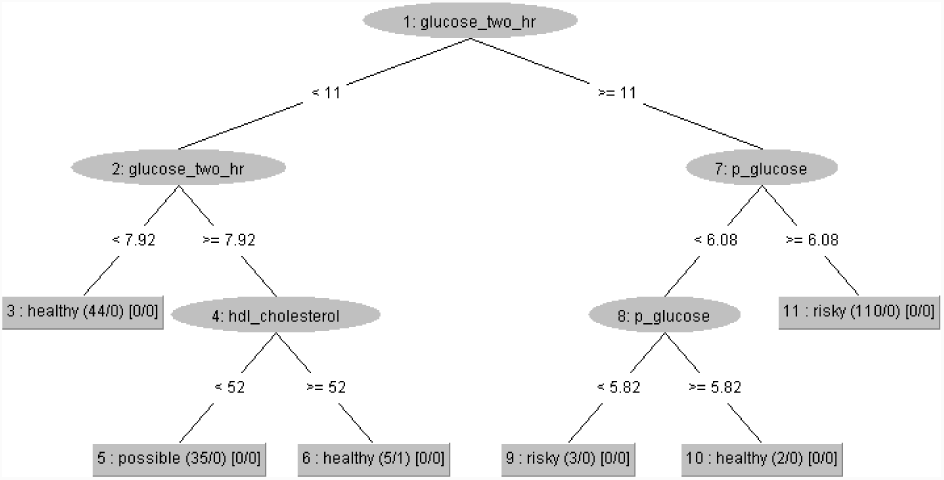
CDT(unpruned) Decision Tree.

From this DT, there are extracted 6 rules where 3 rules show healthy condition, 2 rules represent risky condition and 1 rule explains the possibility of diabetes. From 2, within 50 instances 44 healthy instances are correctly defined by R1 and no instances are misclassified. So, the probability of R1 to describe healthy condition is 100%. R2 also describes 35 instances correctly defined within 36 instances. But for R3, 5 instances are correctly classified but 1 instances are misclassified. On the other hand, R4 fits with 3 instances in risky condition and 2 instances are fitted with R5 without any misclassification. At the end, 110 instances are correctly fitted by R6 without misclassification. So, in this explanation, we can say that R1 is the most frequent rule which detect 44 of 50 instances of healthy condition. Besides, R6 is also most frequent rule which detect 110 instances of 113 for risky condition.

Now, we extract some possible rules from this DT and represent them in table III. From table III, there are generated different rules to predict severity of diabetes. There are remained 13 attributes in the diabetes data, but this model is focused on 3 attributes to represent the severity of diabetes named Plasma Glucose 2hr after glucose (glucose two hr), plasma glucose (p glucose) and HDL-Cholesterol (hdl cholesterol). When glucose two hr is considered less than 11, then it is completely healthy condition for diabetes. But if the range of glucose two hr is considered between greater than/equal 7.46 and less than 11, then it can predict two decision based on hdl cholesterol. In this case, if hdl cholesterol is found less than 52 than it has predicted a possibility about diabetes. But if it is found greater than or equal 52, then it can think healthy condition for people. On the other hand, when When glucose two hr is considered greater than/equal 11 and p glucose is found greater than/equal 6.08, then it is risky condition for diabetes. When glucose two hr is considered between greater than/equal 11 and P glucose, then it depends on p glucose values. In this case, if p glucose is found less than 5.82, then it is risky condition of diabetes. But if it is also found greater than/ equal 5.82, then it is healthy condition.

**TABLE III.**
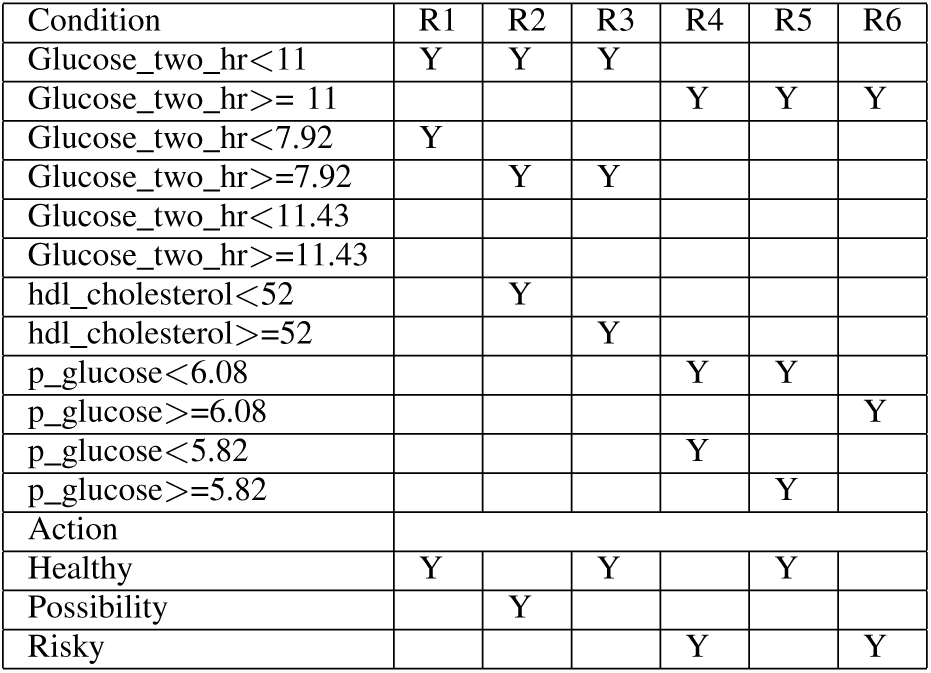
Decision Table of J48 Decision Tree

| Condition | R1 | R2 | R3 | R4 | R5 | R6 |
| --- | --- | --- | --- | --- | --- | --- |
| Glucose_two_hr<11 | Y | Y | Y |  |  |  |
| Glucose_two_hr>= 11 |  |  |  | Y | Y | Y |
| Glucose_two_hr<7.92 | Y |  |  |  |  |  |
| Glucose_two_hr>=7.92 |  | Y | Y |  |  |  |
| Glucose_two_hr<11.43 |  |  |  |  |  |  |
| Glucose_two_hr>=11.43 |  |  |  |  |  |  |
| hdl_cholesterol<52 |  | Y |  |  |  |  |
| hdl_cholesterol>=52 |  |  | Y |  |  |  |
| p_glucose<6.08 |  |  |  | Y | Y |  |
| p_glucose>=6.08 |  |  |  |  |  | Y |
| p_glucose<5.82 |  |  |  | Y |  |  |
| p_glucose>=5.82 |  |  |  |  | Y |  |
| Action |  |  |  |  |  |  |
| Healthy | Y |  | Y |  | Y |  |
| Possibility |  | Y |  |  |  |  |
| Risky |  |  |  | Y |  | Y |

## VI. Conclusion

In this work, we explored significant features to find out the severity of diabetes. Some limitations are considered in this experiment such as 220 records are collected which are small quantity to explore significant rules for predicting diabetes. Besides, we only analyzed diabetes patients of Noakhali except other district’s in Bangladesh. This proposed model is designed in a way that it could be extended and improved for the automation of diabetes analysis. This study may also assist many researcher to optimize possible symptoms of diabetes which is helpful for treatment of this disease in future. In future, we may think about more data with significant attributes that is helpful to find significant attributes in inside and outside in Bangladesh.

## Acknowledgment

We are thankful to Noakhali Diabetes Association for providing diabetes patient data for this research work.

